# Quiescent neural stem cells transiently become ‘neurons’ to coordinate reactivation

**DOI:** 10.1101/2024.11.19.624366

**Authors:** Laura-Yvonne Gherghina, Jocelyn L.Y. Tang, Andrea H. Brand

**Author notes:** Authors contributed equally (Laura-Yvonne Gherghina, Jocelyn L.Y. Tang).

## Abstract

Reactivation of quiescent neural stem cells (qNSCs) is a coordinated process that generates new neurons and glia. Here, we show that during development reactivation follows a hierarchical sequence, whereby anteriorly located qNSCs control the reactivation of more posterior qNSCs in the central nervous system (CNS). Electrical communication between qNSCs and neurons enables this precise spatial and temporal control of reactivation throughout the entire span of the CNS. Using single-cell RNA-seq we discovered that, remarkably, qNSCs switch on a host of neuronal genes while maintaining a stem cell identity. This transient mixed identity is specific to qNSCs, as neuronal genes are turned off after stem cells resume proliferation. Our results uncover long-range communication between qNSCs to coordinate reactivation, enabled by a transient ‘stem cell-neuron’ fate.

## Introduction

Quiescence is an actively maintained state of reversible cell cycle arrest: cells stop proliferating and remain stalled in either the ‘G_0_’ or G_2_ phase of the cell cycle (*1*). Both developing and adult tissues maintain pools of quiescent stem cells (*2*). Cell cycle re-entry (reactivation) of quiescent stem cells occurs during growth and in response to injury or disease, to generate, or replace, differentiated cells. During *Drosophila* development, neural stem cells become quiescent in late embryogenesis and reactivate in early larval development (*3*), giving rise to neurons and glia that contribute to the adult nervous system.

The ability of stem cells to generate new progeny raises the prospect of manipulating stem cells *in vivo* for brain repair or regeneration. To achieve this aim, however, requires knowledge of the signals to which quiescent stem cells respond and an understanding of how the signals are received and acted upon. The efficient and timely reactivation of quiescent neural stem cells is necessary to maintain tissue homeostasis, and for regeneration and repair mechanisms.

In the *Drosophila* CNS, NSCs are located in three major spatial niches along the anterior to posterior axis: brain lobes, thoracic ventral nerve cord (VNC) and abdominal VNC. Upon feeding, NSCs reactivate in response to glia-derived insulin signalling across the CNS (*4, 5*). Once this signal is secreted, it is not known whether the subsequent exit from dormancy is cell-autonomous, or whether long-range non-cell-autonomous signals exist between spatially defined NSC populations in the CNS.

NSCs in *Drosophila* show a dramatic morphological change during quiescence, extending projections into the axon-dense neuropile. Upon reactivation, the NSCs retract these projections and resume asymmetric cell division to produce neurons and glia. The existence of these projection has been known for decades (*3*), yet its function remain elusive. NSCs are distributed across the entire central nervous system, raising the question of how NSC reactivation is coordinated temporally and spatially.

We show that reactivation of qNSCs relies on electrical communication with neurons, which regulates reactivation in a temporally and spatially defined sequence. This enables the coordination of NSC reactivation across the entire central nervous system, from the brain lobes to the nerve cord.

We found that, whilst maintaining the expression of stem cell genes, qNSCs adopt a neuronal morphology and express a host of neuronal genes. NSCs take on a transient mixed ‘stem cell-neuron’ identity that enables communication with neurons to direct reactivation along the length of the CNS. This transient mixed ‘quiescent stem cell-neuron’ identity enables communication with neurons to direct reactivation along the length of the CNS.

## Results

### NSCs reactivation occurs in an anterior to posterior sequence

The majority of NSCs in *Drosophila* enter quiescence late in embryogenesis. In early larval development, feeding induces the secretion of insulin-like peptides from a glial niche, which leads to NSC reactivation from quiescence (*4, 5*). We previously found that 75% of NSCs arrest in the G_2_-phase of the cell cycle, and the remaining 25% arrest in G_0_ (*1*). We showed that heterogeneity in NSC cell cycle arrest leads to differences in reactivation timing, with G_2_-arrested cells reactivating much earlier than their G_0_-arrested counterparts (*1*).

qNSC reactivation initiates in response to feeding, beginning with cell growth (Fig. 1a). We sought to understand how qNSCs coordinate their timing of reactivation. We profiled the distribution of G_2_ and G_0_-arrested cells in the brain lobes and the VNC. We found that the 3:1 ratio of G_2_/G_0_ arrested qNSCs is maintained in the brain lobes and the thoracic VNC, whereas the abdominal NSCs are equally split between G_2_ and G_0_ arrest (Fig. 1b). We assessed reactivation based on expression of the protein Worniu, which is switched on during cell growth at the onset of reactivation, prior to cell division (*1*). We found that qNSCs reactivate first in the brain lobes (4hrs after larval hatching - ALH), followed by qNSCs in the thoracic (8hrs ALH) and abdominal (20hrs ALH) VNC (Fig. 1c and d). We showed previously that G_2_-arrested cells reactivate before G_0_-arrested cells (*1*), but this does not account for the anterior to posterior sequence that begins in the brain lobes (Fig. 1d). Intriguingly, qNSCs in the abdominal VNC remain quiescent more than fifteen hours longer than qNSCs in the brain lobes, despite the synchronous secretion of insulin-like peptides across the entire CNS emanating from the glial-niche (*4*). This suggests that there must be an additional mechanism regulating reactivation.

**Fig 1.**
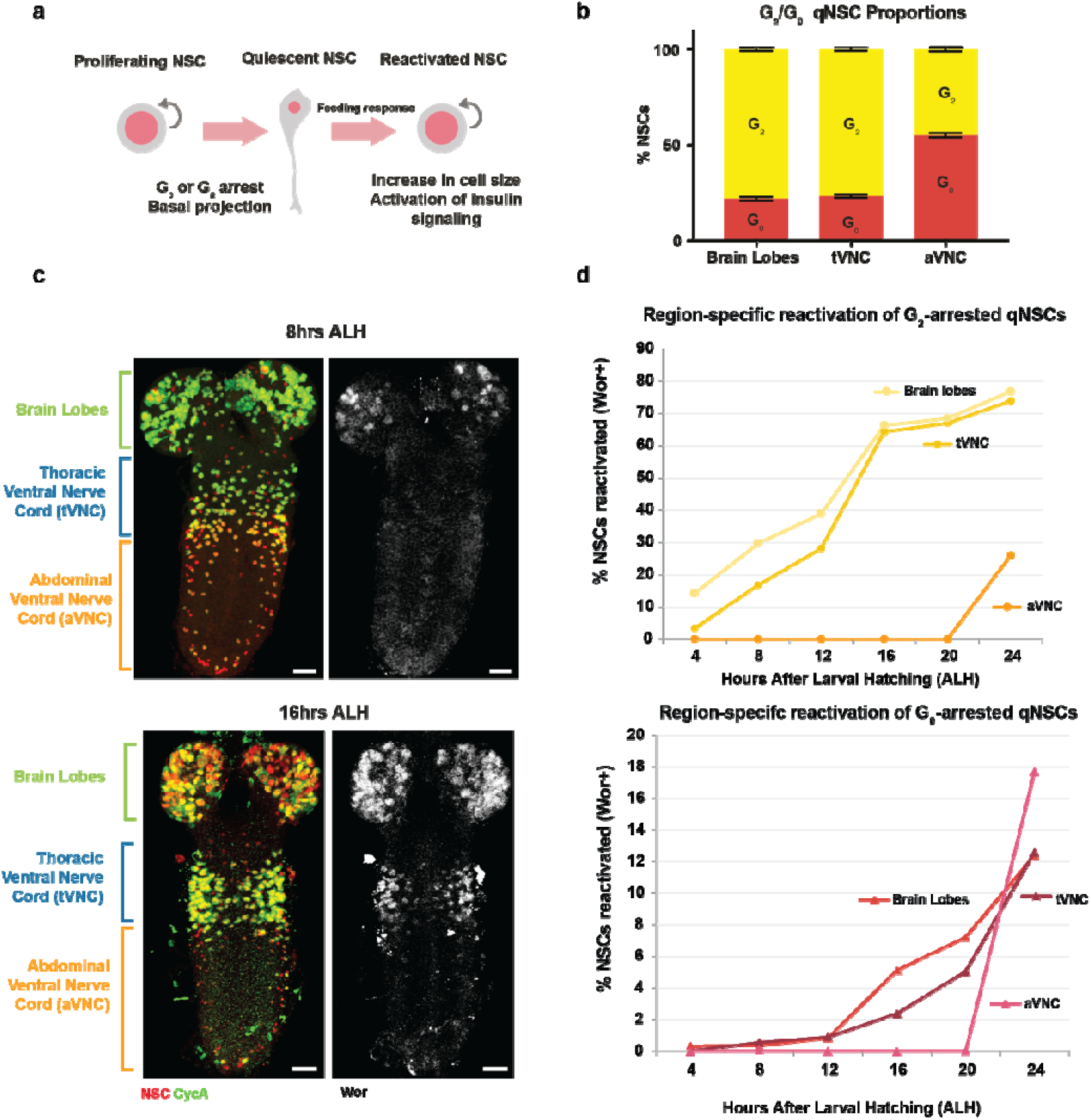
Quiescent NSCs reactivate in an anterior-to-posterior sequence. **a**, Illustration of characteristics specific to quiescent and reactivated NSCs. **b**, Proportion of NSCs arrested in G_2_ or G_0_ in the brain lobes, thoracic VNC (tVNC) and abdominal VNC (aVNC). n=10 brains **c**, Expression of the reactivation indicator Worniu (Wor) in the brain lobes at 8hrs after larval hatching (ALH) followed by tVNC expression at 16hrs ALH; Worniu protein is only expressed in proliferating neural stem cells (Otsuki and Brand, 2018), as an early indicator of reactivation; scale bars, 10um; Deadpan (Dpn, NSC marker) in red, Cyclin A (CycA) in green, Wor in white. **d**, Percentage of NSCs reactivated (Wor+) between 0hrs and 24hrs ALH assayed at 4-hour intervals, split by re-entry from G_2_ or G_0_ arrest; n=4-6 brains.

To achieve this sequence of reactivation, qNSCs may either be able to determine their specific reactivation timings cell-autonomously, or the process could be coordinated via non-cell-autonomous signals from the brain to the VNC.

### Anterior-to-posterior quiescent NSC reactivation is non-cell-autonomous

To determine whether the sequential reactivation of qNSCs follows a cell-autonomous or non-cell-autonomous programme, we generated a genetic tool to restrict expression to NSCs in the brain lobes (Fig. 2a).

**Fig 2.**
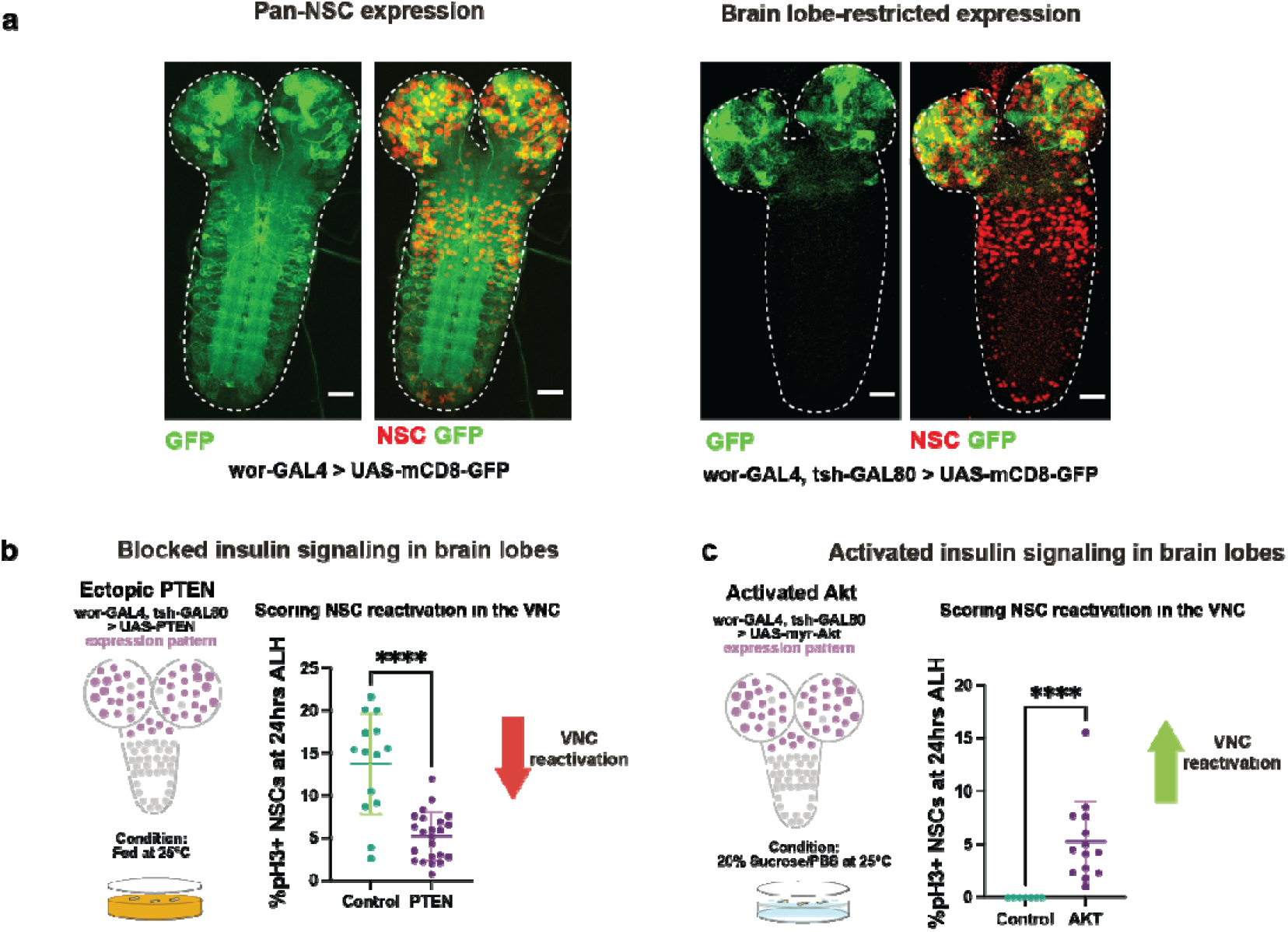
NSC reactivation is non-cell-autonomous. **a**, Ectopic gene expression is restricted to the brain lobes using *wor-GAL4, tsh-GAL80*; Scale bar 10um; mCD8-GFP in green, Dpn in red. **b**, PTEN misexpression in the brain lobes under fed conditions leads to decreased reactivation in the VNC (pH3+ NSCs); n=14 (control), n=23 (PTEN) **c**, Misexpression of constitutively active Akt under amino acid starvation conditions in the brain lobes is sufficient to induce reactivation in the VNC; n=7 (control), n=14 (Akt) Mann-Whitney U-test; ****P<0.0001.

qNSCs reactivate in response to glial-derived insulin-like peptides (*4*). Therefore, we tested first whether inhibiting the insulin signalling pathway in the brain lobes would impair reactivation of qNSCs in the ventral nerve cord. Upon expression of an inhibitor of insulin signalling (PTEN) in the brain lobe qNSCs, we found that reactivation of ventral nerve cord qNSCs was severely impaired (Fig. 2b).

Next, we cultured animals under ‘starvation’ conditions, in which glial insulin secretion is absent and qNSCs cannot reactivate (*4*). We reactivated qNSCs exclusively in the brain lobes by activating insulin signaling. Remarkably, this was sufficient to induce reactivation of the ventral nerve cord qNSCs (Fig. 2c). Therefore, reactivation of NSCs in the brain lobes overcomes the need for glia-derived insulin secretion and is sufficient to induce reactivation. Conversely, even in the presence of glia-derived reactivation signals, preventing reactivation qNSCs in the brain lobe niche dramatically reduces reactivation in the downstream VNC niches. Our results demonstrate that the reactivation signal is propagated between NSCs in a defined sequence from the anterior to the posterior CNS.

Timely reactivation is therefore not only dependent on glial-derived insulin signals, but it also requires a hierarchical sequence from the anteriorly located NSCs to the downstream NSC spatial niches in the VNC. To coordinate reactivation between the brain lobe and VNC niches, NSCs must be able to propagate signals along the entire anterior to posterior axis of the CNS.

### Neuronal activity is required for neural stem cell reactivation

Interestingly, qNSCs extend long cellular projections towards the pre-existing neuropile. These projections are retracted once the cells reactivate and resume proliferation (*6*) (Fig. 3a). When fully extended, the termini of qNSC projections can be seen close to axonal tracts in the CNS. This led us to investigate whether qNSCs in the ventral nerve cord might receive signals directly from the descending tracts of neurons in the brain lobes.

**Fig 3.**
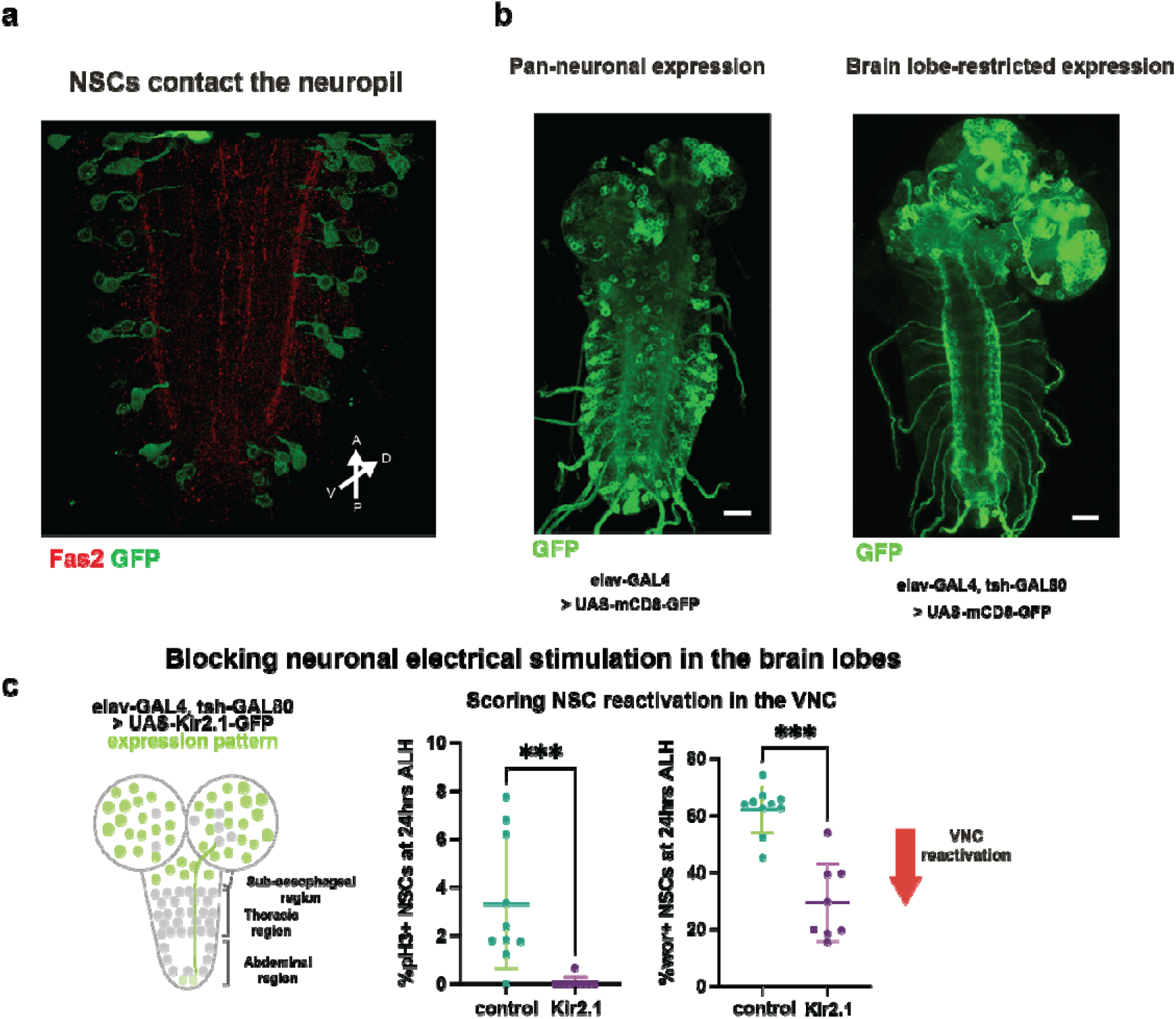
Electrical inactivation of neurons impairs reactivation. **a**, Quiescent NSCs extend a projection towards the neuronal tracts. Maximum intensity projection of the VNC; Fas2 (marking the descending neuronal tracts in the neuropil) in red, *wor*-GAL4 >mCD8-GFP in green. **b**, Ectopic gene expression is restricted to the brain lobes using *elav-GAL4, tsh-GAL80*. Scale bar 10um; mCD8-GFP in green. **c**, Hyperpolarisation of brain lobe neurons and their descending tracts leads to impaired reactivation in the VNC; n=10 (control, pH3), n=9 (Kir2.1, pH3); n=7 (control, Wor), n=10 (Kir2.1, Wor); Mann-Whitney U-test; ***P<0.001.

To test whether neurons in the brain lobes mediate NSC to NSC communication, we specifically silenced brain lobe neurons, which send descending tracts into the VNC, by expressing the potassium channel Kir2.1 ((*7*), Fig. 3b). Blocking neuronal firing drastically delayed the onset of reactivation as marked by Worniu (Fig. 3c) and completely abolished the proliferation of qNSCs in the ventral nerve cord, which was assessed via pH3 staining (Fig. 3c). This demonstrates that neuronal activity is required for the non-autonomous reactivation of posterior qNSCs.

### Quiescent neural stem cells are electrically sensitive

To determine whether qNSCs are able to communicate with neurons in order to propagate signals, we tested whether they were sensitive to electrical activity by hyperpolarising qNSCs using Kir2.1 (Fig. 4a). We found that misexpression of Kir2.1 in the brain lobes led to impaired reactivation of ventral nerve cord qNSCs (Fig. 4b). This suggested that qNSCs are electrically sensitive, which could enable them to communicate with neurons and coordinate their anterior-to-posterior reactivation sequence.

**Fig 4.**
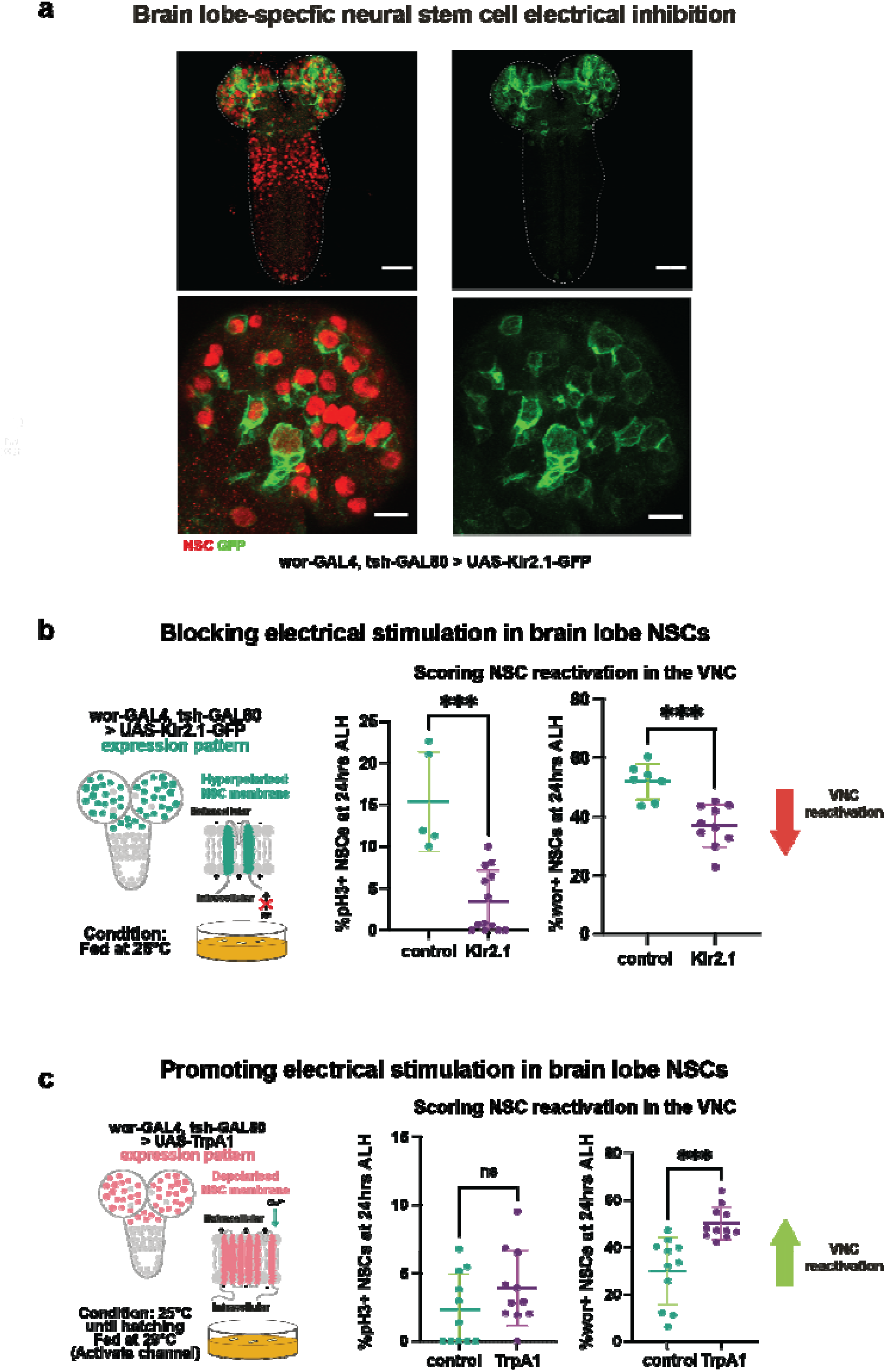
Quiescent NSCs are electrically sensitive. **a**, Misexpression of the potassium channel Kir2.1-GFP in the brain, scale bars, 40um; brain lobe expression of Kir2.1-GFP, scale bars, 10um. **b**, Hyperpolarisation of brain lobe NSCs leads to a delay in reactivation in the VNC; n=5 (control, pH3), n=13 (Kir2.1, pH3); n=7 (control, Wor), n=10 (Kir2.1, Wor). **c**, Depolarisation of brain lobe NSCs using the temperature-sensitive dTrpA1 channel speeds up expression of the reactivation marker Worniu; n=13 (control, pH3/Wor), n=13 (TrpA1, pH3/Wor). Mann-Whitney U-test; ***P<0.001.

Since electrical inhibition of qNSCs prevented reactivation, we tested whether depolarisation, or electrical activation, would enhance cell cycle re-entry. We expressed the temperature-activated Ca^2+^ channel dTrpA1 (*8*) in brain lobe qNSCs and found increased expression of Worniu (Fig. 4c). Therefore, the reactivation of brain lobe NSCs is relayed to NSCs in the VNC and promotes their reactivation. However, there was no significant increase in proliferation, as judged by pH3 labelling (Fig. 4c). This suggests that while depolarisation of the anterior brain lobe qNSCs may accelerate reactivation, it works in concert with other processes.

qNSCs extend long cellular projections toward the axon-rich neuropile. We found that electrical silencing of neurons originating in the brain lobes, which have tracts that descend into the VNC, prevents reactivation in the posteriorly located NSC niches. In addition, electrical silencing of brain lobe qNSCs impairs downstream reactivation, whereas electrical activation accelerates it.

### Quiescent neural stem cells take on the properties of neurons

The morphology of qNSC projections is highly reminiscent of axons. We performed scRNA-seq to identify genes whose expression might regulate qNSC morphology (Fig. 5a).

**Fig 5.**
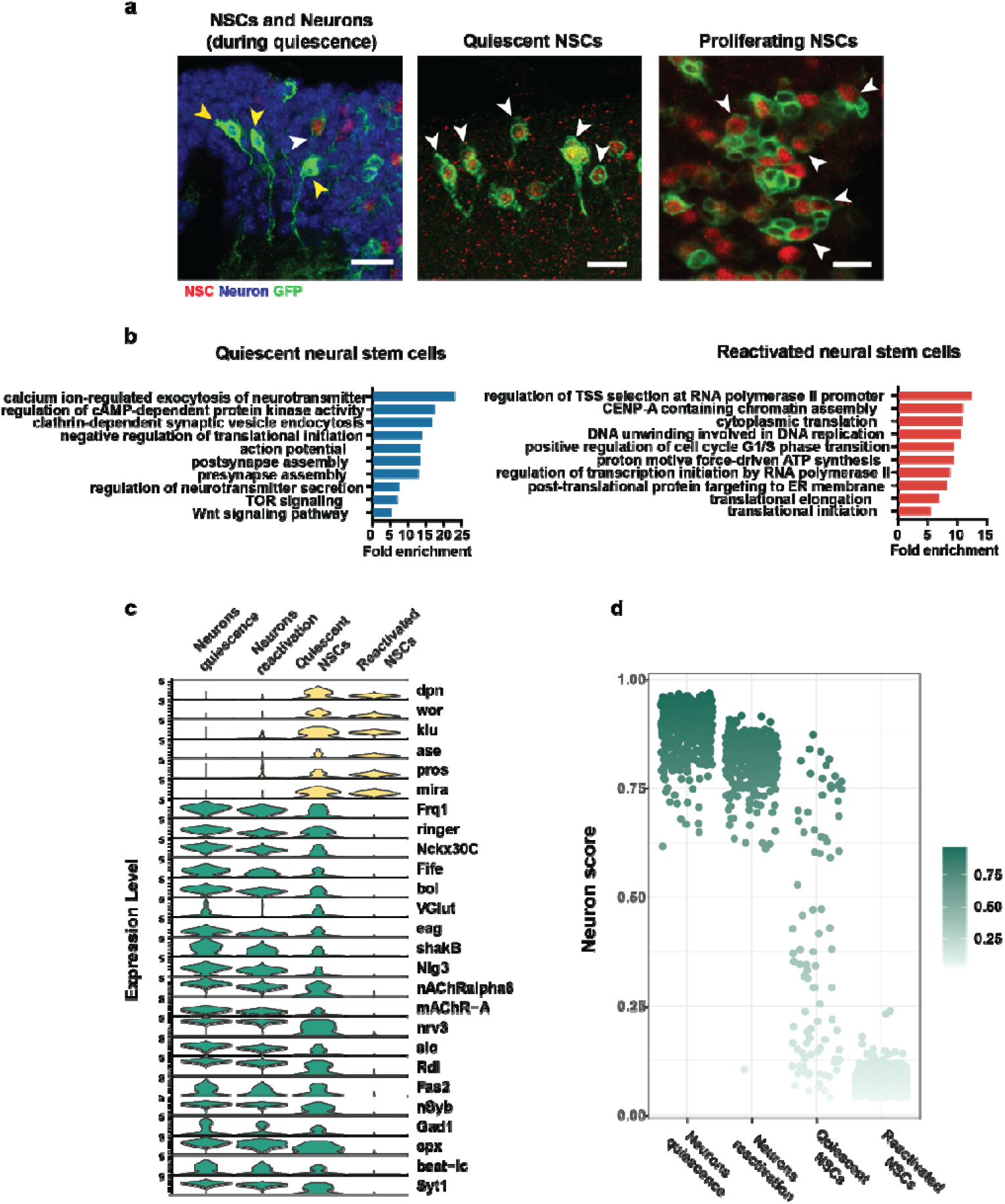
Quiescent NSCs express neuronal genes. **a**, Quiescent NSCs bear a projection that morphologically resembles neuronal axons, which is not present in proliferating NSCs. White arrows indicate NSCs, yellow arrows indicate neurons. **b**, GO terms for quiescent NSCs **c**, GO terms for reactivated NSCs **c**, Expression of NSC and neuronal genes in neurons during quiescence, neurons after reactivation, quiescent NSCs and reactivated NSCs. **d**, Neuron score obtained using UCell comparing neurons and NSCs during quiescence and reactivation.

As expected, we found that both quiescent and reactivated NSCs expressed neural stem cell genes such as *deadpan, worniu* and *klumpfuss*.

Surprisingly, we found that qNSCs also express genes characteristic of neurons (Fig. 5b-d, Supplementary Fig. 1a and b). Neuronal gene expression was only observed in qNSCs; upon reactivation, NSCs reverted to stem cell gene expression and neuronal genes were silenced.

GO term analysis of qNSC gene expression revealed an enrichment for neuronal genes involved in neurotransmitter release, synaptic assembly and synaptic activity (Fig. 5b). In contrast, reactivated NSCs expressed genes for transcription and translation (Fig. 5b). Quiescent, but not reactivated, neural stem cells expressed neuronal genes involved in electrochemical processes, including GABAergic (Gad1, Rdl), cholinergic (nAChRalpha6, mAChR-A) and glutamatergic neurotransmission (VGlut; Fig. 5c). We calculated a “neuron score” based on the top neuronal genes expressed in neurons during late embryogenesis and found that qNSCs have a high neuron score, whereas reactivated NSCs do not exhibit neuronal gene expression (Fig. 5d). Therefore, qNSCs transiently become neuronal while maintaining expression of stem cell genes. This cell fate plasticity allows NSCs to propagate reactivation using electrical communication with neurons along the anterior to posterior axis of the CNS. By adopting a ‘stem cell-neuron’ mixed identity, qNSC reactivation is controlled both temporally and spatially.

## Discussion

Here we show that, in addition to the temporal heterogeneity that exists between the earlier-reactivating G_2_-arrested and later-reactivating G_0_-arrested cells, qNSCs exit dormancy in an anterior to posterior sequence. qNSCs coordinate their spatially and temporally defined sequence of reactivation non-cell-autonomously, beginning in the brain lobes (anteriorly) followed by the ventral nerve cord (posteriorly). We found that reactivation of the posterior stem cells is absolutely dependent upon prior reactivation of brain lobe stem cells, even in the presence of reactivation signals induced by feeding.

Nearly 40 years ago, Truman and Bate first noticed that qNSCs extend cellular projections reminiscent of neurons (*3*), although the functional significance of neural stem cell projections has remained elusive. We show that, remarkably, reactivation occurs through direct communication between neurons and NSCs. We were surprised to discover that qNSCs transiently take on neuronal properties during the quiescence period: they express numerous neuronal genes that are required for communication with neurons, whilst continuing to express neural stem cell genes. In this way, reactivating NSCs in the brain lobes are able to relay signals to qNSCs in the ventral nerve cord via axonal tracts descending from the brain (Fig. 6a and b). These descending axons are contacted by qNSC cellular projections, and communication between the neurons and qNSCs relies on neuronal electrical activity.

**Fig 6.**
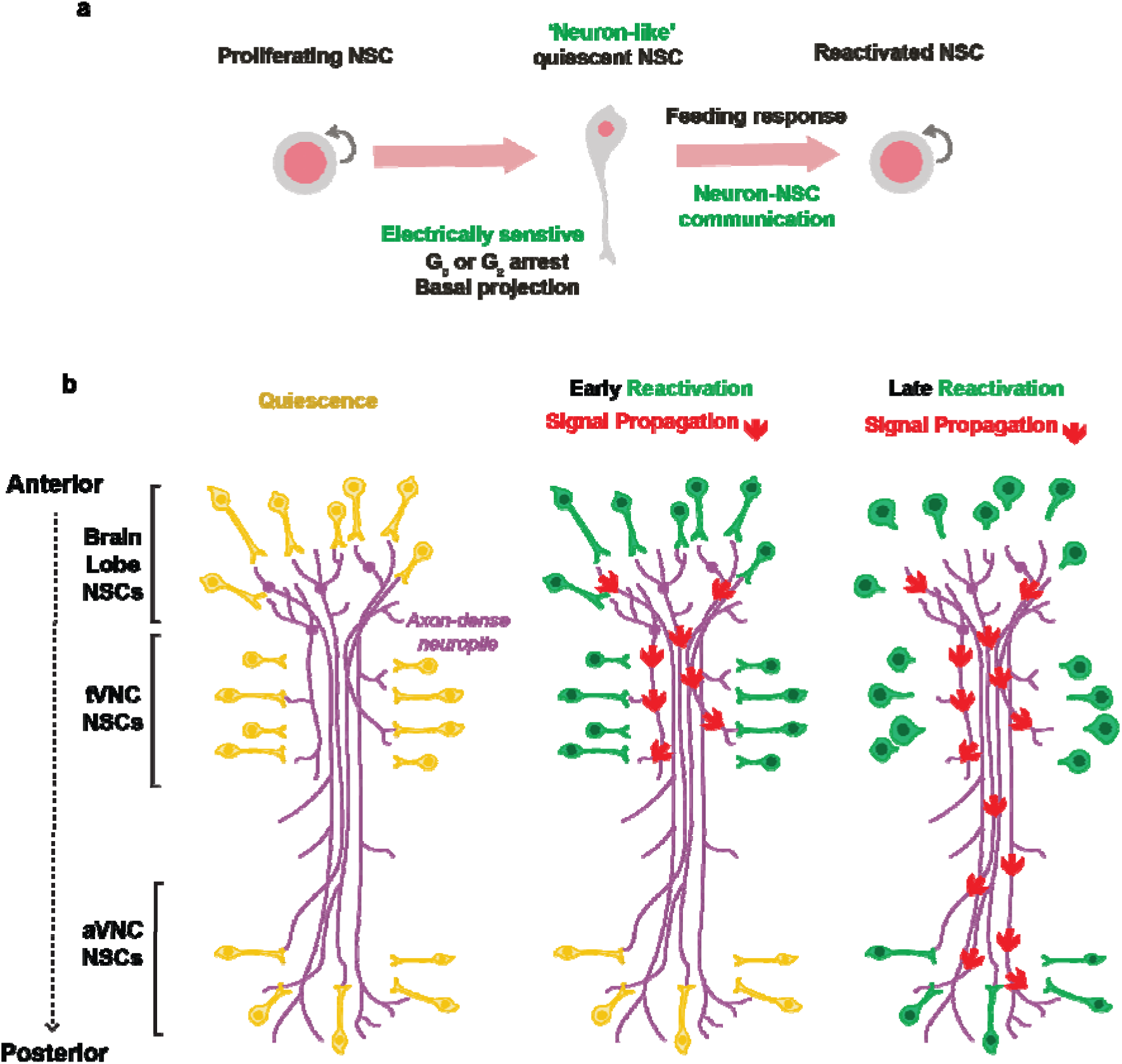
Quiescent NSCs adopt neuronal characteristics to coordinate reactivation via neuronal signalling. **a**, Quiescent NSCs adopt a unique neuronal morphology which allows for communication with neurons within the neuropile. Proliferating NSCs pre- and post-quiescence do not maintain this morphology. Our experiments suggest that the neuronal morphology allows for direct neuron-NSC communication. **b**, NSCs reactivate in a spatially and temporally controlled manner, along the anterior-posterior axis. The descending neurons that project from the brain lobes to the ventral nerve cord enable the state of the NSCs in more anterior regions to be sensed by NSCs in the posterior region.

A crucial role in maintaining and reactivating neural stem cells is played by niche-derived cues(*9*). Previous reports have described neuronal signalling to NSCs in the mouse brain, where neurotransmitter diffusion is thought to act as a regulator of neurogenesis (*10, 11*). In addition, inhibition of GABA release in a subpopulation of interneurons in the SGZ of adult mice was shown to prevent qNSC reactivation (*12*–*14*). Transsynaptic tracing in the adult mouse V-SVZ identified populations of serotonergic neurons that make contacts with subpopulations of adult NSCs, influencing neurogenesis (*15*). Hypothalamic POMC+ neurons, which regulate hunger and satiety, have also been shown to regulate NSC proliferation in the V-SVZ of adult mice (*16*). Despite these instances of communication between NSCs and neurons, no previous study has found that neurons act as mediators of timely reactivation signals between NSCs in distinct spatial niches. In addition, we show that NSC-neuron interactions rely on cell fate plasticity, whereby qNSCs transiently adopt neuronal characteristics. Having neuronal characteristics, whilst maintaining their stem cell fate, enables qNSCs to electrically communicate with neurons in order to coordinate the timing of reactivation in spatially distinct niches along the CNS. Future work will explore the specific neuronal circuits that coordinate stem cell reactivation within the anterior to posterior hierarchical sequence.

Neuron-NSC interactions may also be relevant for cancer progression and pathology. Glioma cells have been observed to form excitatory synapses with neurons in the surrounding microenvironment (*17, 18*). Electrochemical signals have been shown to influence the proliferation of glioma cells, which form synapse-like structures with glutamatergic neurons (*17*). In paediatric gliomas, depolarisation of glioma tumour cells via optogenetics promotes proliferation (*19*). In some cases, the formation of synapses between glioma cells and neurons also impacts the frequency of seizures, a common symptom in glioma patients (*20*). Although the identification of neuronal-like glioma cells has been well observed (*17*), the specific cell identity of these cells has not yet been identified, in light of the cellular heterogeneity identified by single-cell transcriptomics (*21*). It is currently unknown whether all cell types in glioma have the ability to form synapses. Our finding that quiescent NSCs can adopt neuronal characteristics may be relevant in the disease context since some glioma cells are also quiescent (*22*) and developmental mechanisms are aberrantly activated in tumourigenesis.

Our study has identified a non-cell-autonomous spatial hierarchy by which quiescent NSCs coordinate timely reactivation. This mechanism is mediated via NSC-neuron communication, achieved through the transient acquisition of neuronal characteristics by quiescent NSCs. Further investigation of specific neuronal circuits and potential role of neurotransmitters will shed light on NSC-neuron communication during brain development, extending towards how these mechanisms may be involved in disease.

## Materials and Methods

### Fly husbandry and stocks

Embryos were collected on apple juice plates supplemented with wet yeast. Larvae were fed with standard sugar/yeast agar (SYA) food and kept at 25°C. For amino acid starvation experiments, larvae were moved into 20% sucrose in PBS.

The following lines were used: wor-GAL4(*23*), elav-GAL4(*24*), UAS-mCD8-GFP (*25*), tubGAL80^ts^ (*26*), tsh-GAL80 (gift from Gero Miesenboeck), UAS-myr-AKT (*27*), UAS-PTEN (Huang et al., 1999), UAS-Kir2.1-GFP (*7*), UAS-TrpA1 (BL# 26263), w^1118^.

### Immunochemistry

Larval brains were dissected in PBS and fixed in 4% formaldehyde in PBS for 20 minutes at RT. Dissected tissues were washed in 0.3% Triton in PBS (PBST) for 3 × 5 minutes at RT. The following primary antibodies were used: chicken anti-GFP (1:2000, abcam), guinea pig anti-Dpn(*28*) (1:5000, rabbit anti-pH3 (1:200, Millipore), rat anti-Wor (1:200, abcam), mouse anti-Fas2 (1:100, DSHB), rabbit anti-CycA (1:10,000, gift from David Glover). Antibodies were diluted in 0.3% PBST and samples were incubated overnight at 4°C. Samples were washed 3 × 15 minutes in 0/3% PBST at RT. Secondary antibodies (Alexa Fluor 405, 488, 546, 568, 633 (Life Technologies); DyLight 405 (Jackson Laboratories)) were diluted in PBST at 1:200 concentration and incubated overnight at 4°C. Samples were mounted using Vectashield (Vector Laboratories).

### Imaging and Quantification

Images were acquired using a Leica SP8 upright confocal and a Nikon AX Inverted Confocal. Image processing and cell counting were performed using Fiji (v. 2.14.0). Fig.s were made using Adobe Illustrator (v. 27.2) and graphs were made using Prism (v. 9.5.1). Where applicable, statistical analysis was also performed using Prism. 3D reconstruction was performed using the Leica LASX 3D Viewer software using maximum intensity projection images.

### scRNAseq sample preparation and data analysis

The scRNA-seq dataset used in this study was prepared and analysed in Tang and Brand (in preparation). In brief, stage 17 embryos (quiescence) and 24h ALH (reactivated) larval brains were dissociated via mechanical and enzymatic dissociation. In order to enrich for neural cells, the neural-specific driver *wor*-GAL4 was used to express a membrane-bound GFP (UAS-mCD8-GFP) to facilitate fluorescence-activated cell sorting (FACS) on single-cell suspensions. Live, (Propidum iodide-negative) and GFP+ cells were isolated using a S800Z Cell Sorter (Sony). Cells were then processed using the Chromium Next GEM Single Cell Library and Gel Bead Kit (v2 and v3.1) (10X Genomics). Sequencing was performed on a Novaseq 6000 (SP well) (Illumina) by the CRUK Genomis Facility.

FASTQ sequencing files were de-multiplexed and processed using CellRanger v7.1.10 (10X Genomics) and output files were analysed using Seurat V4 (*29*). Cells with less than 200 and more than 6000 unique mRNA feature counts, more than 5% mitochondrial counts were excluded from analysis. Resulting dataset were analysed with default parameters as described the Satija lab (satijalab.org).

We used the UCell package(*30*) in R to generate a neuronal score based on the top 30 genes expressed in the neuronal cluster during quiescence (stage 17). GO term enrichment analysis was performed using PANTHER 17.0 (*31*).

## Acknowledgements

We would like to thank the Gurdon Institute Imaging Facility (GIIF) for technical support. We thank Leia Judge and Anna Malkowska for help with preparation of scRNA-seq samples. We also would like to thank Katarzynia Kania at the Cancer Research UK Cambridge Institute Genomics Core Facility for sample preparation and sequencing of samples.

## Author contributions

L.Y.G, J.L.Y.T and A.H.B designed the experiments. L.Y.G and J.L.Y.T performed the experiments. L.Y.G, J.L.Y.T and A.H.B analysed the data and wrote the manuscript.

---

**Fig Supplementary 1.**
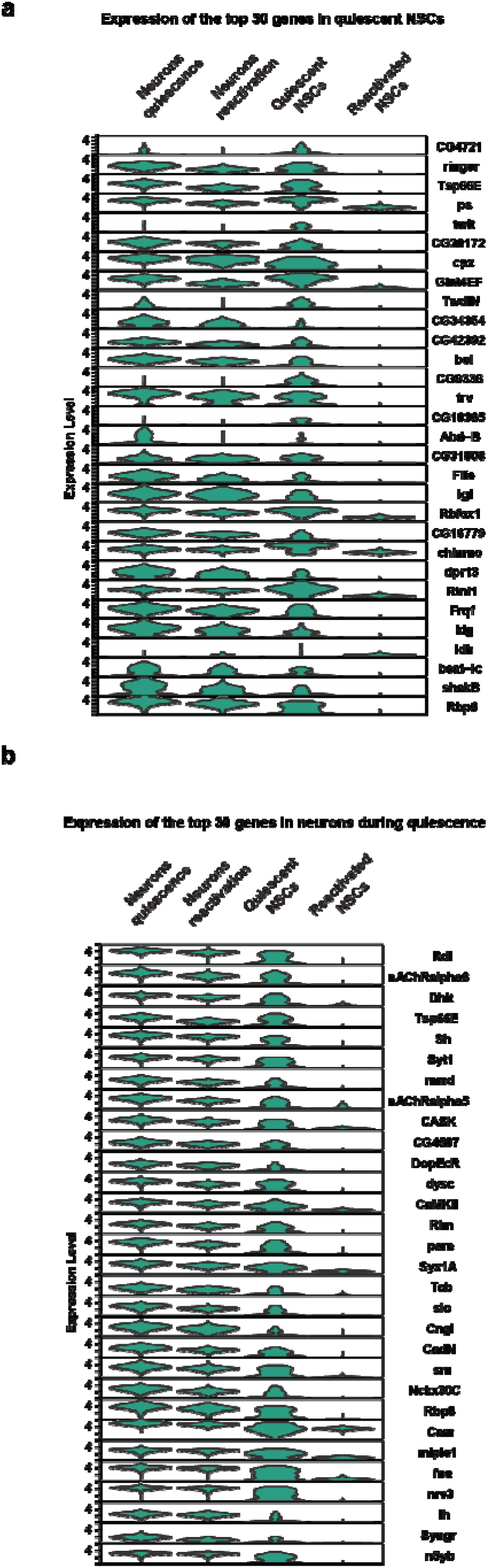
Expression of genes in NSCs from single-cell transcriptomic data. **a**, Violin plots showing the average expression of the top 30 genes enriched in qNSCs. Expression levels of these 30 genes are shown in neurons (while NSCs are quiescent), qNSCs, neurons (while NSCs are reactivating) and reactivating NSCs. **b**, Violin plots showing the average expression of the top 30 genes enriched in neurons. Expression levels of these 30 genes are shown in neurons (while NSCs are quiescent), qNSCs, neurons (while NSCs are reactivating) and reactivating NSCs.

